# Gene Expression Profiles Reveal Potential Targets for Breast Cancer Diagnosis and Treatment

**DOI:** 10.1101/2022.09.03.504469

**Authors:** Mohammad Hossein Nasirpour, Mohammad Sabery Anvar, Nasirpour Alireza, Salimi Mahdieh, Sepahyar Soheil, Minuchehr Zarrin

**Affiliations:** Department of Biology, Islamic Azad University Science and Research Branch, Department of Medical Genetics, Institute of Medical Biotechnology, National Institute of Genetic Engineering and Biotechnology (NIGEB), Tehran, Iran; Department of Systems Biotechnology, National Institute of Genetic Engineering & Biotechnology (NIGEB), Iran; Department of Electrical and Computer Engineering, Batten College of Engineering, Old Dominion University, Norfolk, VA, United States; Department of Medical Genetics, Institute of Medical Biotechnology, National Institute of Genetic Engineering and Biotechnology (NIGEB), Tehran, Iran; Department of Computer Sciences, Michigan Technological University, United States

## Abstract

Figuring out the molecular mechanisms underlying breast cancer is essential for the diagnosis and treatment of this invasive disorder. Hence it is important to identify the most significant genes correlated with molecular events and to study their interactions in order to identify breast cancer mechanisms. Here we focus on the gene expression profiles, which we have detected in breast cancer. High-throughput genomic innovations such as microarray have helped us understand the complex dynamics of multisystem diseases such as diabetes and cancer. We performed an analysis using microarray datasets by the Networkanalyst bioinformatics tool, based on a random effect model (REM). We achieved pivotal differential expressed genes like *ADAMTS5, SCARA5, IGSF10*, and *C2orf40* that had the most down-regulation, and also *COL10A1, COL11A1*, and *UHRF1* that they had the most up-regulation in four-stage of breast cancer. We used CentiScape and AllegroMCODE plugins in CytoScape software in order to figure out hub genes in the protein-protein interactions network. Besides, we utilized DAVID online software to find involved biological pathways and Gene ontology, also used Expression2kinase software in order to find upstream regulatory transcription factors and kinases. In conclusion, we have found that the statistical network inference approach is useful in gene prioritization and is capable of contributing to practical network signature discovery and providing insights into the mechanisms relevant to the disease. Our research has also identified novel transcription factors, kinases, pathways, and genes that may serve as important targets for the development of diagnostic biomarkers and treatments.

## Introduction

Breast cancer is a significant public health concern, as it is the most commonly diagnosed cancer and ranks second in the population of women as a cause of death. Outbreaks in most countries are growing, given recent attempts to prevent the disease[1]. This is because breast cancer is a complicated disease with several contributing factors influencing disease progression. Despite several kinds of research, neither the exact etiology of breast cancer nor the mechanisms behind the patient--heterogeneity, are identified. For this, breast cancer diagnosis and treatment remain both a daunting and fascinating task[2]. With the rapid expansion of genome-wide gene expression profiling techniques, a range of bioinformatics data analysis frameworks were established to classify genes linked to breast cancer and to uncover gene identities for prognosis and estimation of care. Because breast cancer is a complex disease, nevertheless, it should not only be ascertained by individual genes but also through the organized impact of multiple genes[3].

The classification of breast cancer is based on histological characteristics, the appearance of cancerous cells and the presence of specific biomarkers like epidermal growth factor receptor (ErbB2/Her2), progesterone receptor (PR), and estrogen receptor (ER) [4, 5]. ERα correlated with signaling pathways, active in breast cancer, and inhibition of it would either decrease breast cancer occurrence or has a healing effect on early breast cancer. [6]. Membrane estrogen receptor could initiate various signaling pathways such as PI3K/Akt/mTOR and MAPK, and the former is one of the significant pathways, engaged in cancer development and induced by different receptor tyrosine kinases like IGF-1, EGFR, and FGFR, while inhibited by phosphatase and tensin homolog (PTEN). From that, mutation or reduction of PTEN expression would be associated with breast cancer occurrence [7]. The mutation in *TP53*, *PIK3CA*, *GATA3*, *Map3K1*, *MLL3*, and *CDH1* are the most noticeable ones in breast cancer [8]. Also, MAPK probably triggers unrestrained cell proliferation, which all make chemo and radiotherapy ineffective [7].

In addition, in recent decades there has been a trend to approach tumor cells from a molecular and genetic point of view, which has dramatically improved the quality of care, especially with regard to the identification of one particular gene whose expression is altered in the genetic context of one particular form of cancer. The advent of precision medicine facilitates tumor profiling concerning genetic and genomic aspects would benefit the detection of variable types of breast cancer to appoint proper medication [8]. There are also few novel genomic assays (Oncotype DX, MammaPrint, and PAM50) from which a prognostic assessment of the risk of relapse in breast cancer can be performed at its early stage. The design of the gene expression involved will essentially decide the extent of cancer malignancy, which potentially circumvents the traditional approach to cancer that can be prevented [9].

Furthermore, systems biology, computer-based analyses such as gene set enrichment, differential gene expression, and networking revolutionized breast cancer study in terms of identification of involved biomarker and signaling pathway which would improve understanding of disease complexity [10, 11]. Besides, researchers have extensively applied the technique of meta-analysis to study networking, which would be particularly applicable for more investigation into transcriptional data integration. Several bioinformatics methods, such as PINTA7, prioritize genes by integrating data on gene expression with the protein-protein interaction (PPI) network through a random walk approach to enrich and ultimately re-rank the candidate genes [12].

Although breast cancer has been conventionally subtyped, molecular techniques like microarray could benefit classification of breast cancer in terms of involved genes and proteins in biological processes [13]. Improvement in genomic profiling such as different types of sequencing techniques will reduce the adverse effect of missing clinical data, i.e. it is feasible to compare the genomic or proteomic environment of breast cancer in the sense of clinical studies and in the direction of advancing in bioinformatics tools, cancer diagnosis and prediction of its response to therapy would be considerable [8].

We conduct the transcriptomic analysis with two qualified microarray datasets to classify DEGs in four stages of breast ductal carcinoma tissues to obtain a more accurate prediction of genetic history variations involved in this malignancy. These DEGs performed clustering research and PPI network. The findings could guide us in finding new biomarkers and therapeutic targets.

## Methods

### Microarray datasets and Preprocessing

For the first step of our analysis, we tried to collect transcriptomic datasets for breast cancer and corresponding normal cells, originated from the Gene Expression Omnibus (GEO) data- base (https://www.ncbi.nlm.nih.gov/geo/|), based on some indications like tissue type, cancer stage (not grade) on Jun 31, 2017 using the following query:

“Breast cancer” [Title] OR “breast tumor” [Title] OR “breast carcinoma” [Title] AND “Homo sapiens” [Organism] AND “expression profiling by array” [Filter] AND “normal” NOT “therapy” NOT “drug” NOT “treatment” AND “stage I and IV”.

Stages I, II, III and IV were selected based on the clinical records that accompanied each sample.

After removing samples either produced using just grading or treated cells or taken from peripheral blood or containing siRNAs, we concluded with two independent GEO datasets with accession numbers of GSE53752 and GSE61304. In this study, our focus was about stages of breast cancer that were classified by clinical history. The GSE53752 raw data was already normalized by robust multi-array averaging (RMA) algorithm and GSE61304 which we did normalized by publicly available expression console software v 1.3.1 which is specific for Affymetrix dataset [14], using the RMA algorithm[15]. Probe sets summarization files (CHP) were generated for each gene. As a part of this project, probe sets were entered into Geworkbench 2.5.1 package[16] and non-useful data were filtered based on two criteria: omission of data without Entrez identification and recognition markers of multiple probe set. Probe sets with the highest mean expression value were retained.

### Identification of DEGs and visualization

Based on meta-analysis of microarray datasets, differentially expressed (DE) genes can be defined. We used Networkanalyst (http://www.networkanalyst.ca/faces/home.xhtml), which is a web interface for integrative meta-analysis of the microarray[17]. Combat procedures using empirical bios methods were used for elimination of the batch effect [18]. Annotation of the datasets was done by translating the gene symbols into respective Entrez IDs after importing the data sets into software. The quantile normalization of the data was used and the data sets were reviewed for data integrity before further proceeding[19]. We used combined effect size and Cochran’s Q test for statistical analysis, and selected Random Effects Model (REM) method with a significance level of P < 0.05 to combine P values from the multiple datasets for meta-analysis [20]. Differential expression analysis was performed with Networkanalyst for each dataset, independently using a false discovery rate of 0.05, a significance of P < 0.05, and Limma algorithm-based t-test (LAT).

Networkanalyst efficiently support comprehensive evaluation of differential gene expression and constructed network. We chose STRING interactome[21] (https://string-db.org/) with high confidence score and cut-off of 900 (High confidence) to visualize and assess network interaction between proteins. Besides, we chose First-order Network, made of seed nodes and their direct interactions with neighboring ones to remove the machine error during optimization of Network Analyst.

### Identification of sub-networks

The proteins were clustered inside the network to detect the location of the densely populated area. This was achieved by Cytoscape [22] plugins including the algorithm of Allegro MCODE plugin that is Molecular Complex Detection (MCODE), based on the weight of nodes and density of neighboring ones. This algorithm performs in three steps Vertex weighting, molecular complex prediction, and post-processing to filter or add protein to the preplanned system [23]. MCODE parameters encompassed degree cutoff, node score, K- score and Max depth of 2, 0.5, 5 and 100.

### Centrality analysis and hub genes identification

Hub genes in core network were detected, using CentiScape plug-in in Cytoscape v 3.5.1. CentiScape computes some major parameters such as betweenness, closeness, and degree of centrality for detected leading nodes, individually, within network role. The betweenness corresponds to the most direct connections pass through the interesting node. The closeness indicates the reciprocal mean distance mean from one spot to other accessible ones. The degree of centrality refers to the total number of neighboring nodes, cooperated with a distinctive node.

### Functional Annotation Clustering of DE Genes

DAVID online webserver (Databases for Annotation, Visualization and Integrated Discovery)[24] was used to identify affected cellular pathways and processes by browsing the DE genes obtained from Networkanalyst webserver and clustered DE genes achieved from CytoScape. The altered functional clusters were rated according to DAVID’s returned enrichment ratings. Enrichment scores above 1.3 (P-value < 0.05) were considered highly significant.

### Up-stream regulatory protein-protein interactions

We used the open source program Expression2Kinas (X2K) which was built on Java platform (java 6 SDK) to determine the upstream regulatory protein-protein interaction network elements. The X2 K is available on the windows, Mac DOS X and Linux platform at’’ http://www.maayanlab.net / X2K”[25]. X2K site contains several new transcription factor libraries and kinase, and PPI networks. Suggested that the X2K pipeline implements a new logical method for defining and rating upstream regulators such as transcription factors (TFs) responsible for the observed shift. Transcription factors and kinases that most likely control observed changes in mRNA expression were prioritized. In the X2K app, we uploaded DEGs obtained from analysis through Networkanalyst. As a result of X2K program outputs and the upstream DEG regulators, all TFs and Kinases were collected.

## Results

Two studies met the inclusion criteria and were chosen for our analysis. All of them provided metadata of high quality which allowed for analysis. In Total, 137 samples were collected, of which 29 were normal and 108 were patient (Table 1). The tumors were staged into, stage I having 17 cases (15.74%), stage II having 58 cases (53.70%), stage III having 29 cases (26.85%) and stage IV having 4 cases (3.70%) were available for this study. In stage four, the number of samples was not adequate that can be attributed to high late-stage patient mortality, and it is important that further attention should be given to stage four for further investigations. Moreover, we used Geworkbench software for data filtration, and from 54675 probesets, 21103 unique probsets with specific Entrez gene ID were achieved.

**Table1:** characteristics of extracted datasets for meta-analysis: The microarray data sets from Gene Expression Omnibus dataset (GEO) which is the one of the international public repositories in the context of high-throughput data; Normal: normal samples; stage: Breast Cancer patients.

| Accession Number | Total Sample | Normal | Stage 1 | Stage 2 | Stage 3 | Stage4 | Chip Type |
| --- | --- | --- | --- | --- | --- | --- | --- |
| GSE61304 | 61 | 4 | 5 | 33 | 18 | 1 | [HG-U133_Plus_2]<br>Affymetrix Human Genome U133 Plus 2.0 Array |
| GSE53752 | 76 | 25 | 12 | 25 | 11 | 3 | Agilent-012097<br>Human 1A<br>Microarray (V2)<br>G4110B |

### Analysis of microarray datasets and DEGs determination for each stage

After quality control and normalization, the expression profile was developed, and a meta-analysis was conducted for each data collection. We performed the data integration to increase data quality [26, 27], increase the statistical calculation [28], and decrease systemic errors [29–31]. Networkanalyst web server have used for data processing and due to the different applied platform, it was preferred to do random effect model choice for analysis of datasets. One of the aims of this study was to identify specific DEGs for each stage. Differentially expressed genes were divided into Up- or Down- regulated genes using our criteria. The P value < 0.05 and a fold-change ≥ 2 were set as the cut off values of DEgenes. Following the above process for each stage related to cancer, the results were shown in table 2, including stages, the overall number of differential expression genes (DEGs), up or down-regulation level (Supplementary file S1-S4). Moreover, the most up-regulation and least down-regulation of genes showed in Table3.

**Table2:** Combination of two datasets and normalization all differentially expressed genes that were obtained with P value < 0.05 and log FC >±2

| Stage | DEGs | UP | Down |
| --- | --- | --- | --- |
| 1 | 4349 | 2555 | 1794 |
| 2 | 5094 | 3066 | 2028 |
| 3 | 3719 | 2088 | 1631 |
| 4 | 2461 | 1157 | 1304 |

**Table 3:** The important up and down regulated differentially expressed genes with p-value `P < 0.05 in four stages of breast cancer`.

| Differentially Expressed Genes |  |  |  |  |  |  |  |  |  |  |  |
| --- | --- | --- | --- | --- | --- | --- | --- | --- | --- | --- | --- |
| down regulated genes |  |  |  |  |  |  |  |  |  |  |  |
| Stage1 | CombinedES | Pval | Stage2 | CombinedES | Pval | Stage3 | CombinedES | Pval | Stage4 | CombinedES | Pval |
| ADAMTS5 | -5.4838 | 3.36E-12 | SCARA5 | -6.0423 | 0 | IGSF10 | -4.8269 | 0.01264 | C2orf40 | -8.4881 | 5.28E-08 |
| TSLP | -5.1017 | 0.014472 | ATP1A2 | -5.346 | 0.000333 | SCARA5 | -4.5417 | 0.017851 | TSLP | -7.6163 | 8.17E-08 |
| EDNRB | -4.8265 | 0.001496 | SDPR | -5.0459 | 0.043781 | TSLP | -4.3937 | 6.09E-08 | MATN2 | -7.5664 | 8.17E-08 |
| LRFN5 | -4.7784 | 0.027934 | TMEM132C | -5.0442 | 0.030054 | LRFN5 | -4.3311 | 0.002292 | MAOB | -6.8609 | 0.002336 |
| IGSF10 | -4.4634 | 4.01E-05 | RERGL | -4.8228 | 0 | HLF | -4.3118 | 6.38E-11 | KLHL29 | -6.1592 | 8.86E-07 |
| PPAP2B | -4.4489 | 3.36E-11 | EDNRB | -4.6097 | 6.71E-05 | PAMR1 | -4.2665 | 8.52E-08 | FAM189A2 | -6.0684 | 8.71E-07 |
| CXCL12 | -4.3849 | 3.56E-11 | GDF10 | -4.6052 | 0 | AK5 | -4.1431 | 1.16E-06 | EDNRB | -5.9676 | 9.29E-07 |
| IL33 | -4.3678 | 0.009774 | OXTR | -4.5902 | 1.25E-06 | PPAP2B | -4.1143 | 9.34E-08 | CRYBG3 | -5.5397 | 1.53E-06 |
| FAM13A | -4.3203 | 3.56E-11 | TIMP4 | -4.5433 | 1.08E-09 | ADAMTS5 | -4.1075 | 0.016989 | TGFBR3 | -5.4953 | 2.07E-06 |
| KL | -4.2894 | 0.00817 | FREM1 | -4.5408 | 0 | SPRY2 | -4.0851 | 8.12E-11 | GALNTL1 | -5.4765 | 2.01E-06 |
| up regulated genes |  |  |  |  |  |  |  |  |  |  |  |
| Stage1 | CombinedES | Pval | Stage2 | CombinedES | Pval | Stage3 | CombinedES | Pval | Stage4 | CombinedES | Pval |
| COL10A1 | 5.1058 | 3.36E-12 | COL11A1 | 4.0578 | 0.001765 | UHRF1 | 4.098 | 0.042025 | COL11A1 | 11.304 | 9.08E-09 |
| MMP11 | 4.7161 | 0.002072 | COL10A1 | 4.0072 | 0.002633 | RRM2 | 3.98 | 7.67E-05 | FUT3 | 10.336 | 9.28E-09 |
| CSTF2 | 4.0785 | 6.35E-11 | MMP11 | 3.8318 | 0 | HN1 | 3.9561 | 8.12E-11 | MMP1 | 6.6762 | 0.002278 |
| HMGB3P1 | 3.81 | 1.29E-10 | WISP1 | 3.7832 | 4.00E-06 | BIRC5 | 3.9024 | 7.59E-05 | C1orf106 | 6.4936 | 3.78E-07 |
| HMGB3 | 3.6927 | 0.003978 | DDX39A | 3.6622 | 0 | LRRC59 | 3.8889 | 8.94E-11 | SYNDIG1 | 6.2233 | 0.029835 |
| UHRF1 | 3.6019 | 0.000652 | RRM2 | 3.6054 | 0 | SPAG5 | 3.7294 | 0.000521 | SULF1 | 5.7826 | 1.54E-06 |
| TK1 | 3.5562 | 3.98E-10 | UHRF1 | 3.5777 | 0 | DLGAP5 | 3.6473 | 0.034465 | INHBA | 5.6078 | 1.53E-06 |
| CCDC167 | 3.5523 | 4.94E-10 | TACC3 | 3.5356 | 0 | MMP11 | 3.5514 | 2.92E-09 | WISP1 | 5.5634 | 1.53E-06 |
| S100A11 | 3.4169 | 8.38E-10 | UBE2T | 3.4741 | 0.01003 | ANLN | 3.4963 | 2.58E-05 | HIST1H1B | 5.4433 | 0.005786 |
| RRM2 | 3.3999 | 7.67E-10 | TK1 | 3.4038 | 9.41E-07 | TACC3 | 3.4683 | 4.25E-10 | PGK1 | 5.3123 | 2.01E-06 |

After comprehensive analysis using DAVID webserver, biological pathways of the most top-ten downregulated genes included in table 3 that are, response to peptide hormone, positive regulation of penile erection, ATP metabolic process, response to drug, organ regeneration, response to cytokine, adult locomotory behavior, regulation of blood pressure, BMP signaling pathway. While, pathways associated with up-regulated genes were collagen catabolic process, negative regulation of B cell differentiation, positive regulation of cellular protein metabolic process, chondrocyte development, endodermal cell differentiation, collagen fibril organization, cell-cell adhesion, protein auto ubiquitination, mRNA 3’-end processing, cartilage development, termination of RNA polymerase II transcription, cell division, hematopoietic progenitor cell differentiation, cell proliferation, microtubule cytoskeleton organization.

At the final part of this process, this software provides the possibility of constructing a network of interference between all DEG(s) in each uploaded dataset. The biggest created network between DEGs in each stage was downloaded from NetworkAnalyst software for a more complete investigation. In addition, we determined number of specific DEGs for each stage by Venn diagram built-in http://bioinformatics.psb.ugent.be/webtools/Venn/. (The complete results of Venn diagram for each stage is in supplementary S5). Following uploading tables, designated for each stage, the different probabilities for share genes have been determined in different stages and each one, separately (Figure1). For stage one 612, stage two 887, stage three 477, and stage four 642 DEGs have recognized, of course, for each stage specifically. Yet, we have considered the most top-ten up and down regulated of them to better analysis and determination of biomarker values.

**Figure1:**
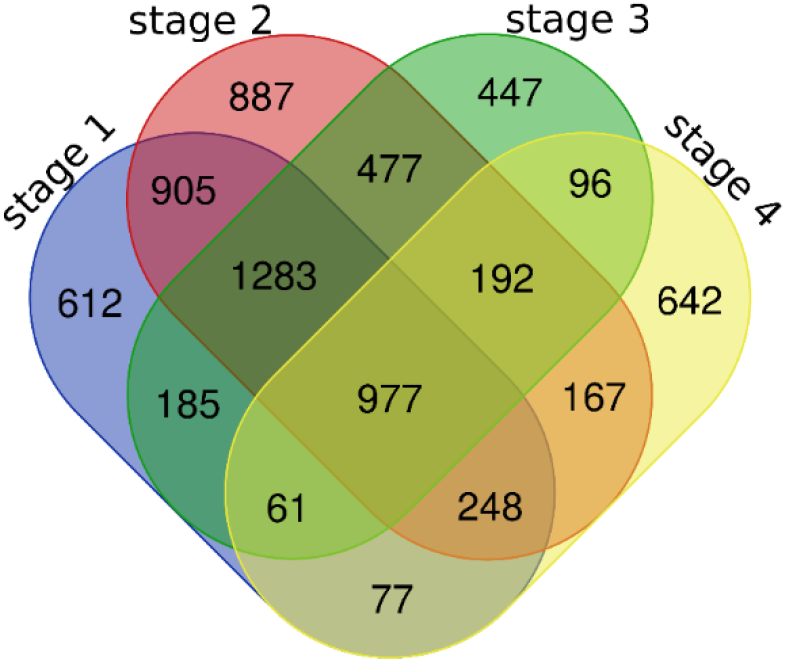
The result of Venn diagram shows specific DEGs for each stage.

### Checking centrality indexes and Gene ontology

Another aim of this study was to introduce pathways involved in different stages of breast cancer. Thus, the constructed networks were clustered based on topological peculiarity. The features of proteins and pathways, in which they are involved, were determined by peculiarity of their corresponding gene ontology through DAVID webserver. (Table4- the results of GO for each stage and cluster). After identification of GO features on each cluster at different stages, betweenness and closeness indices were measured to determine the key or hub protein available on those clusters. From that, the important nodes on each cluster were ascertained (table 4).

**Table4:**
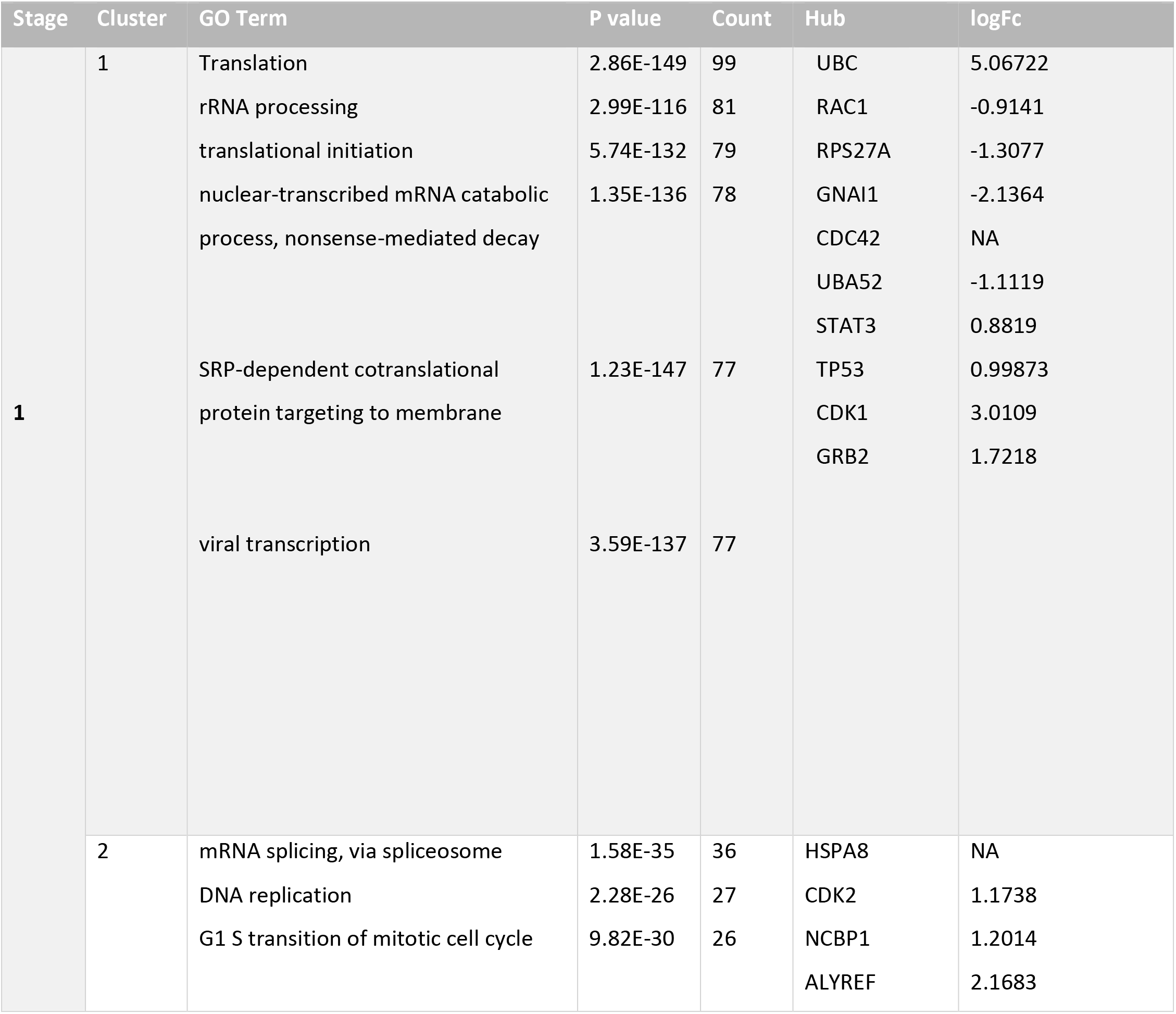
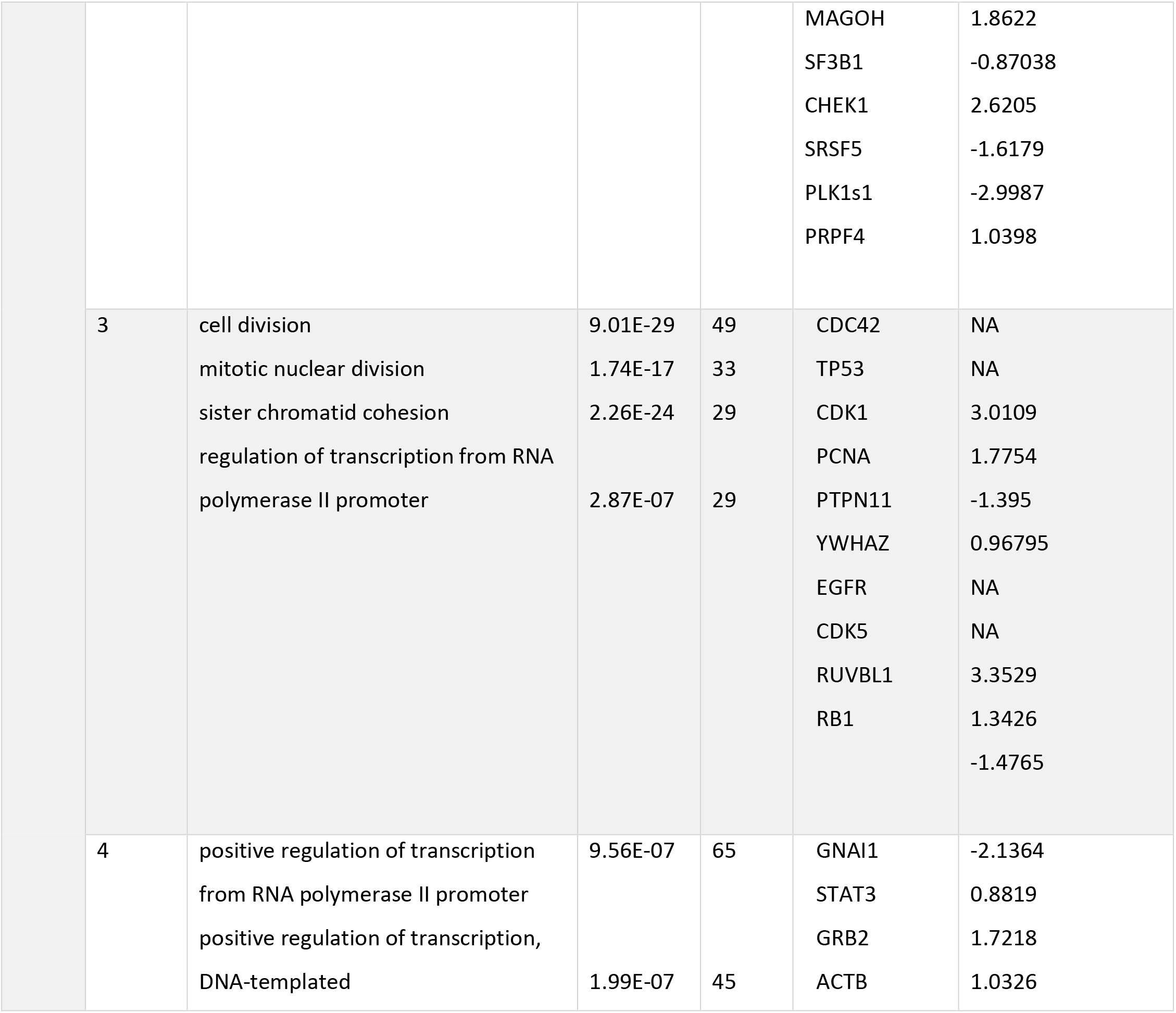
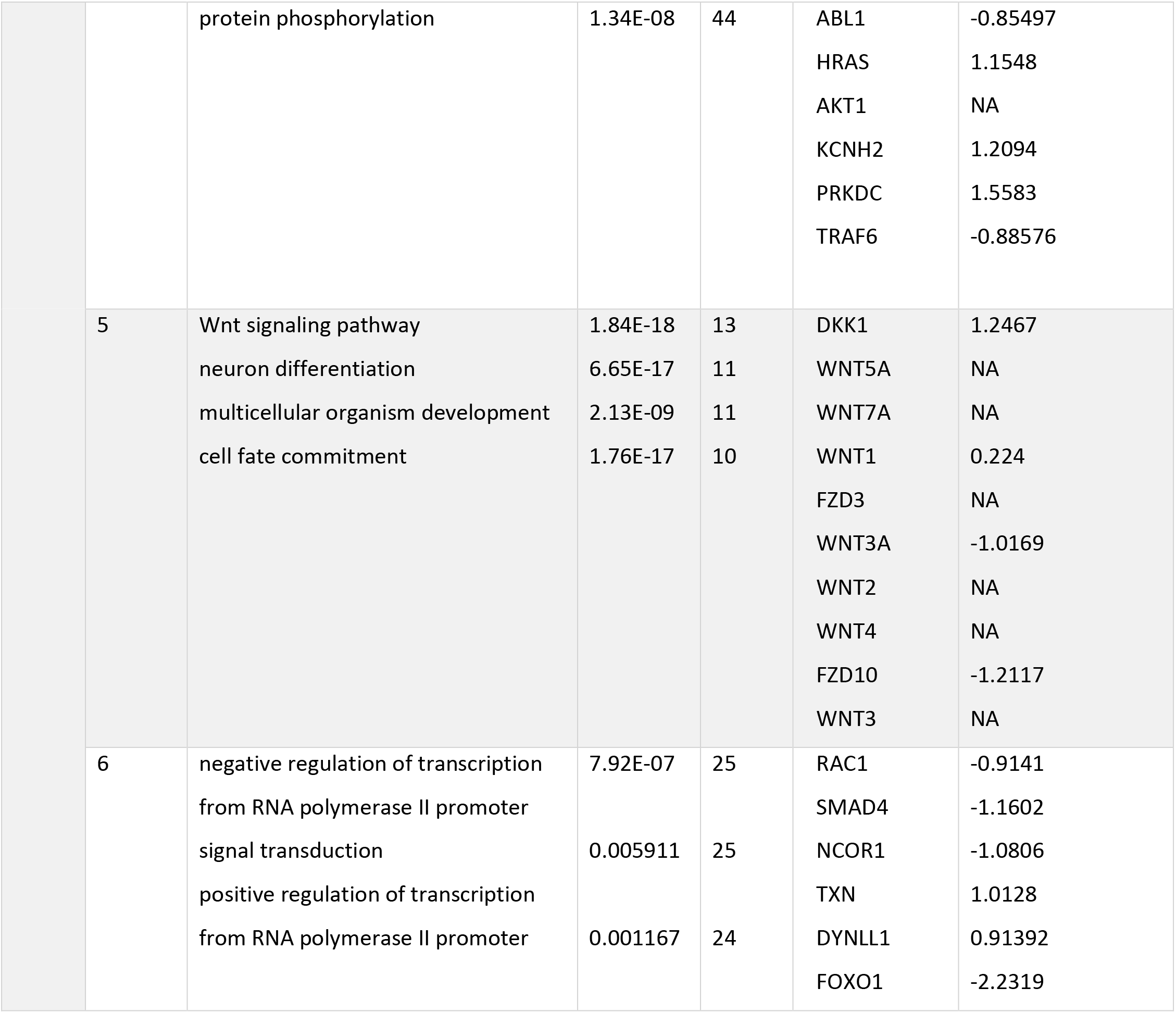
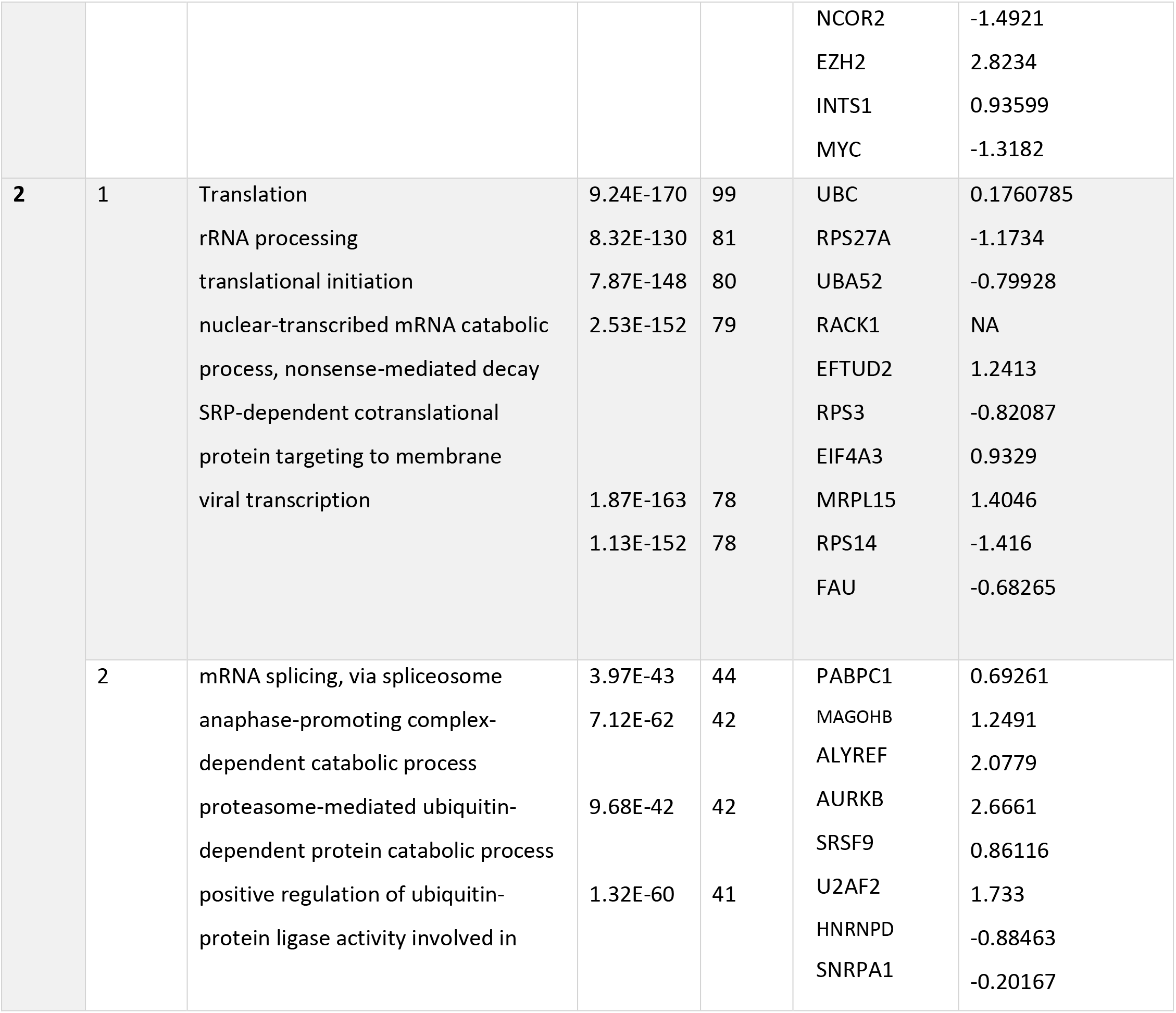
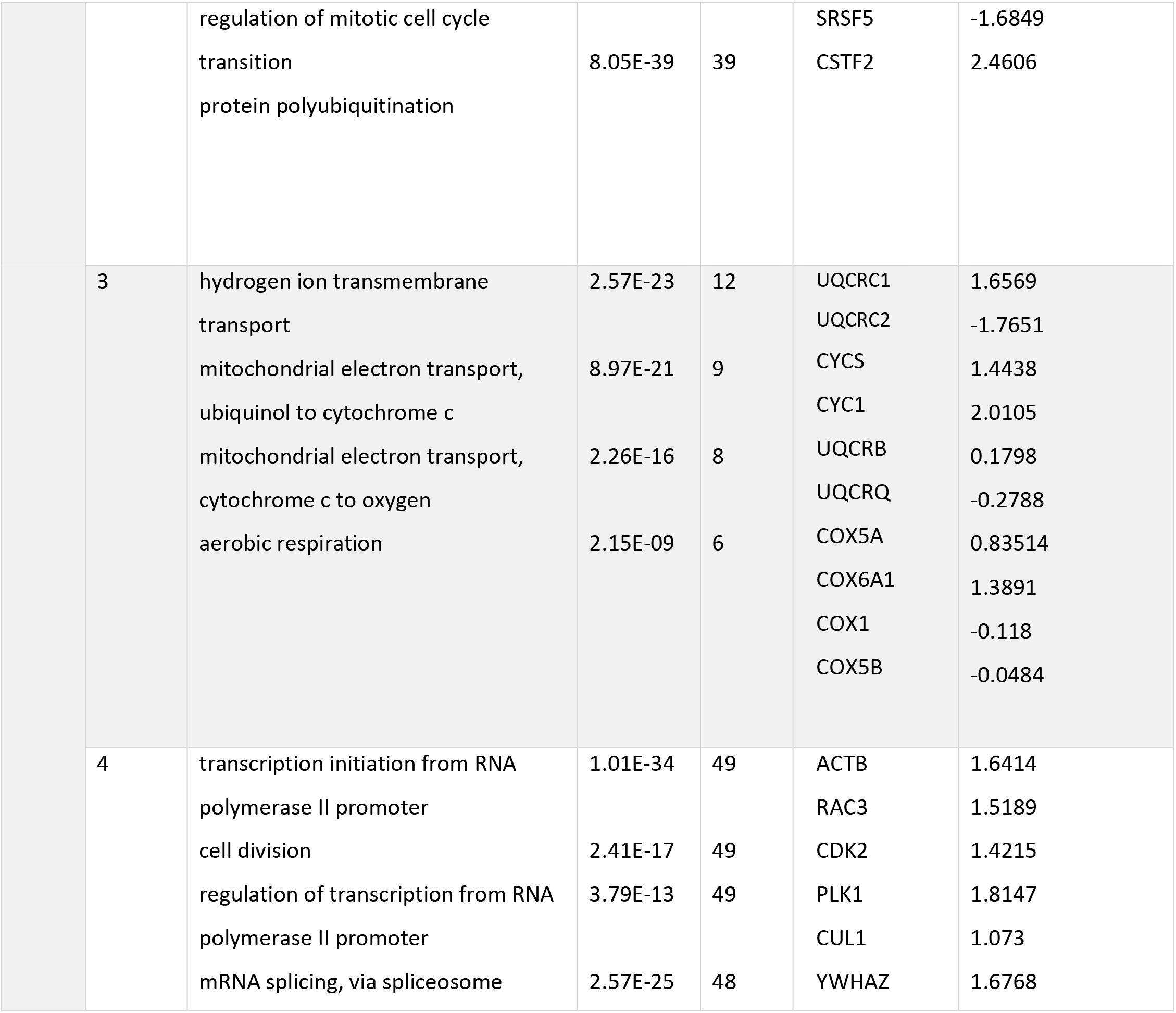
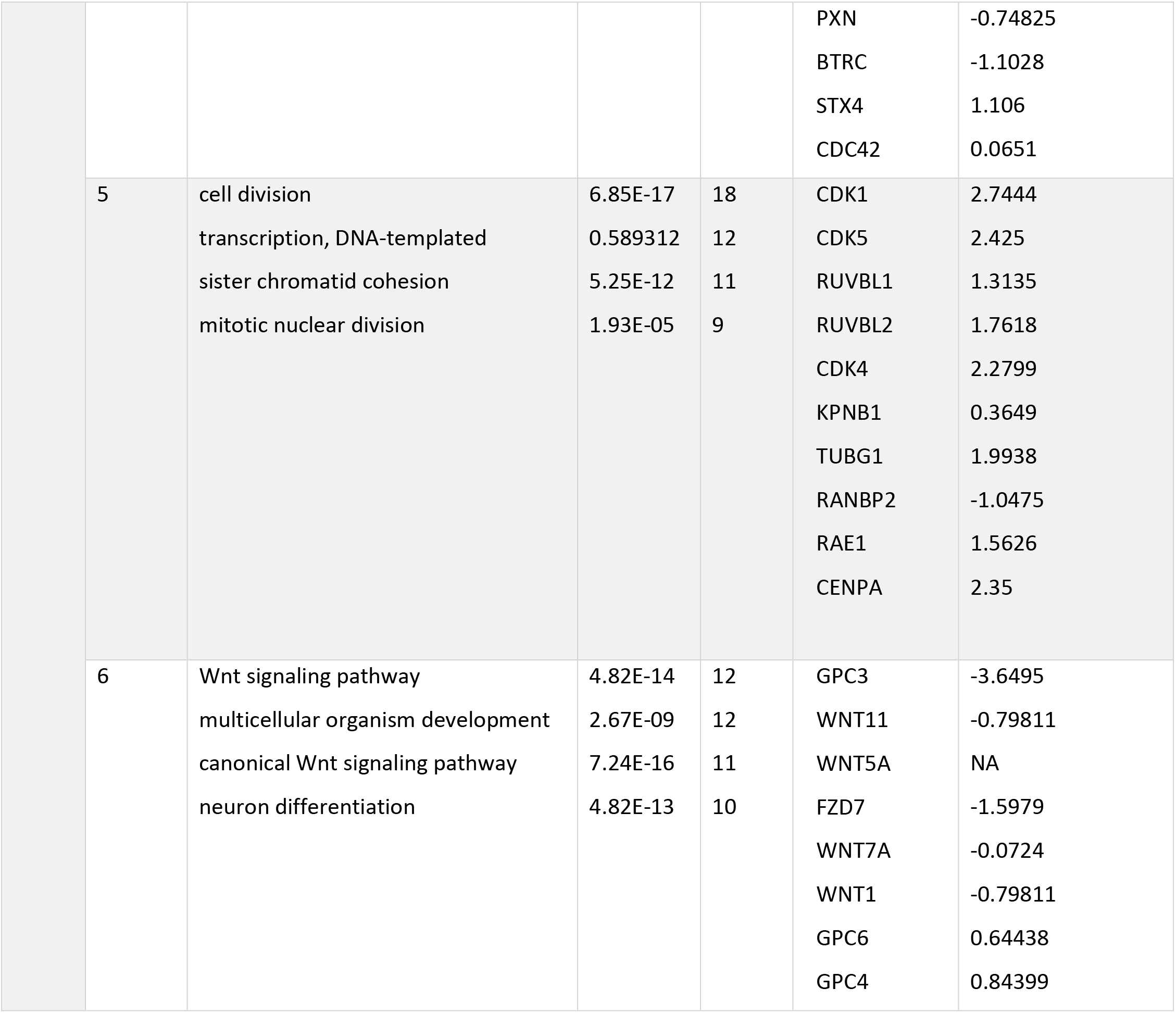
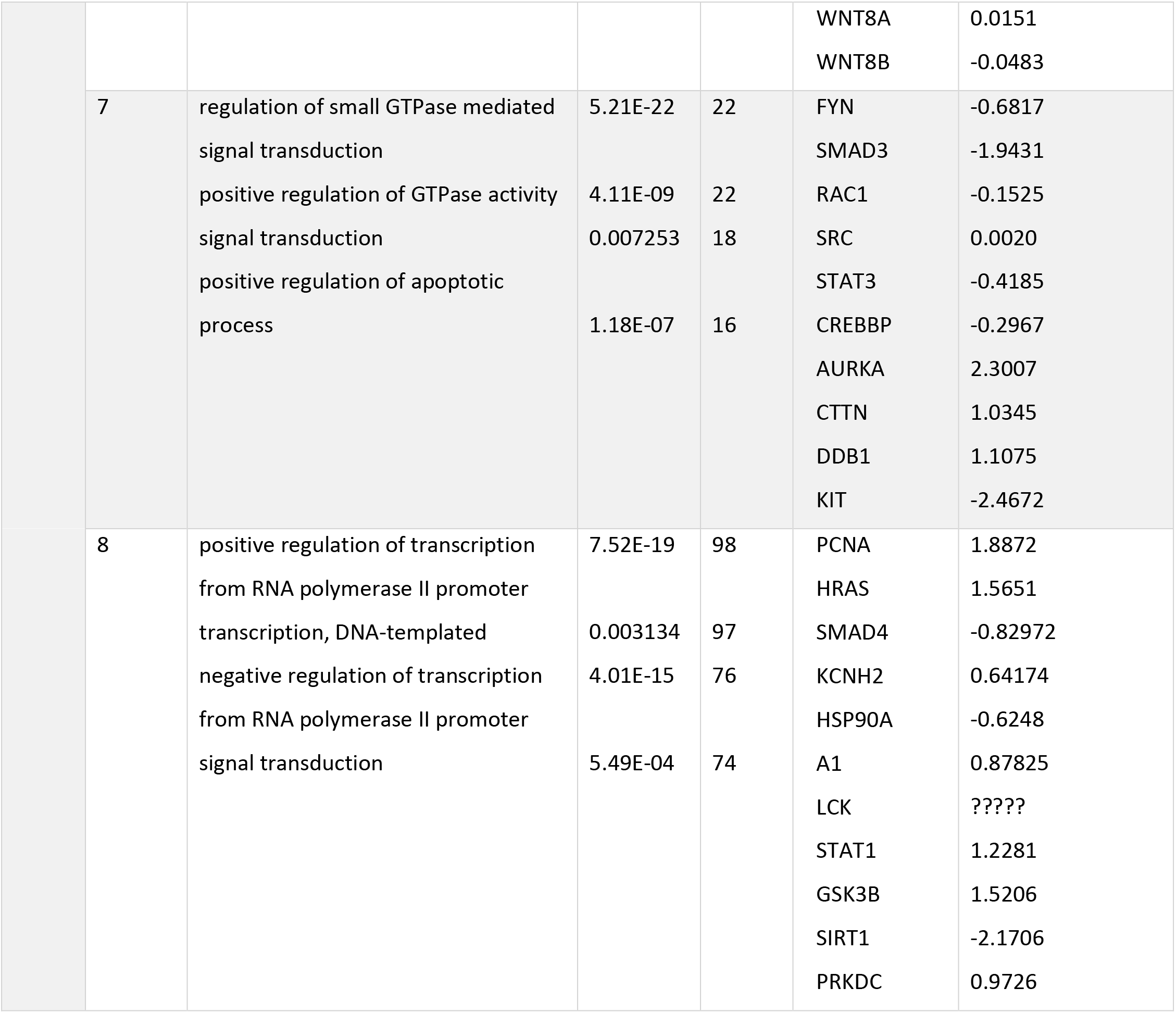
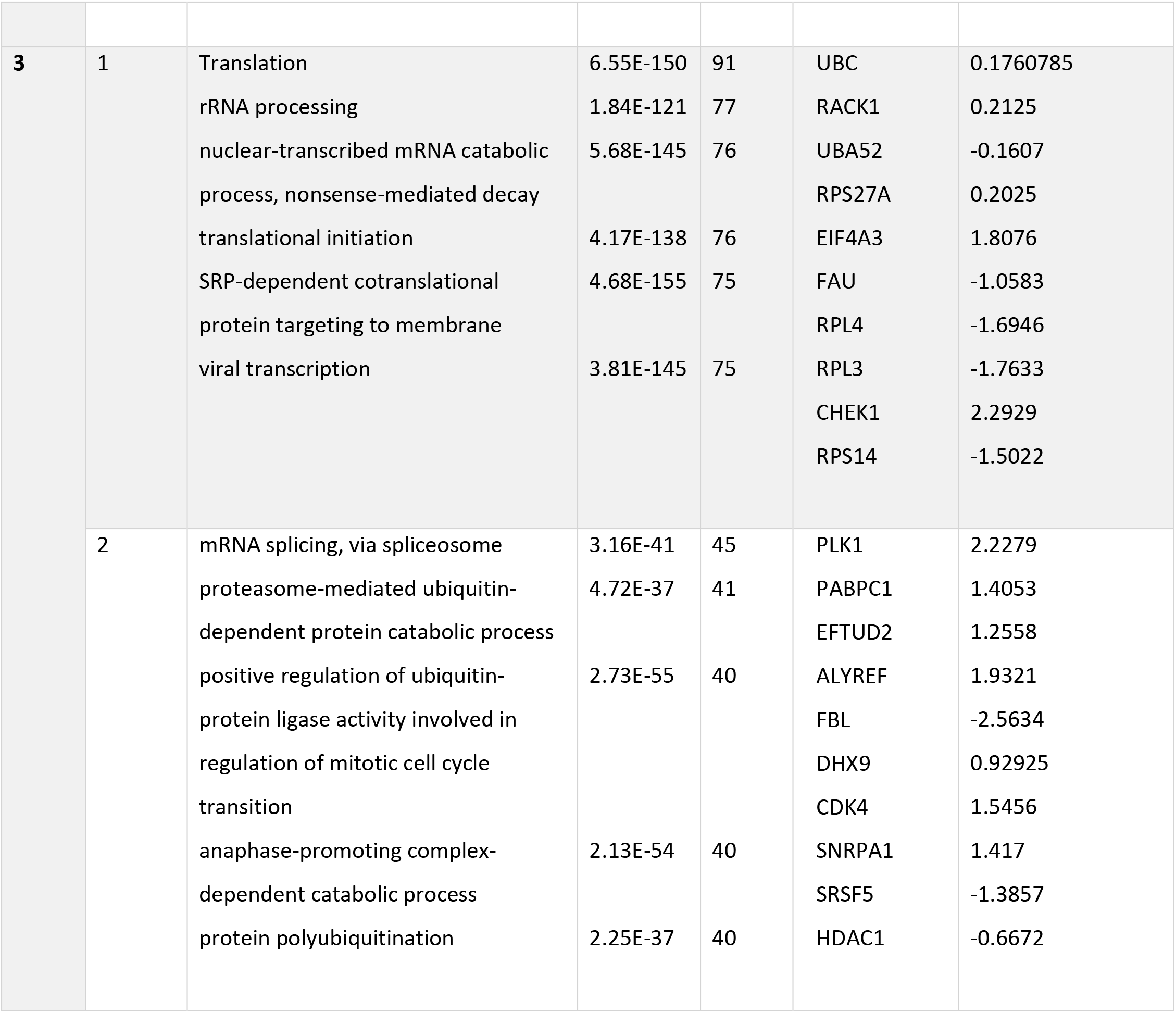
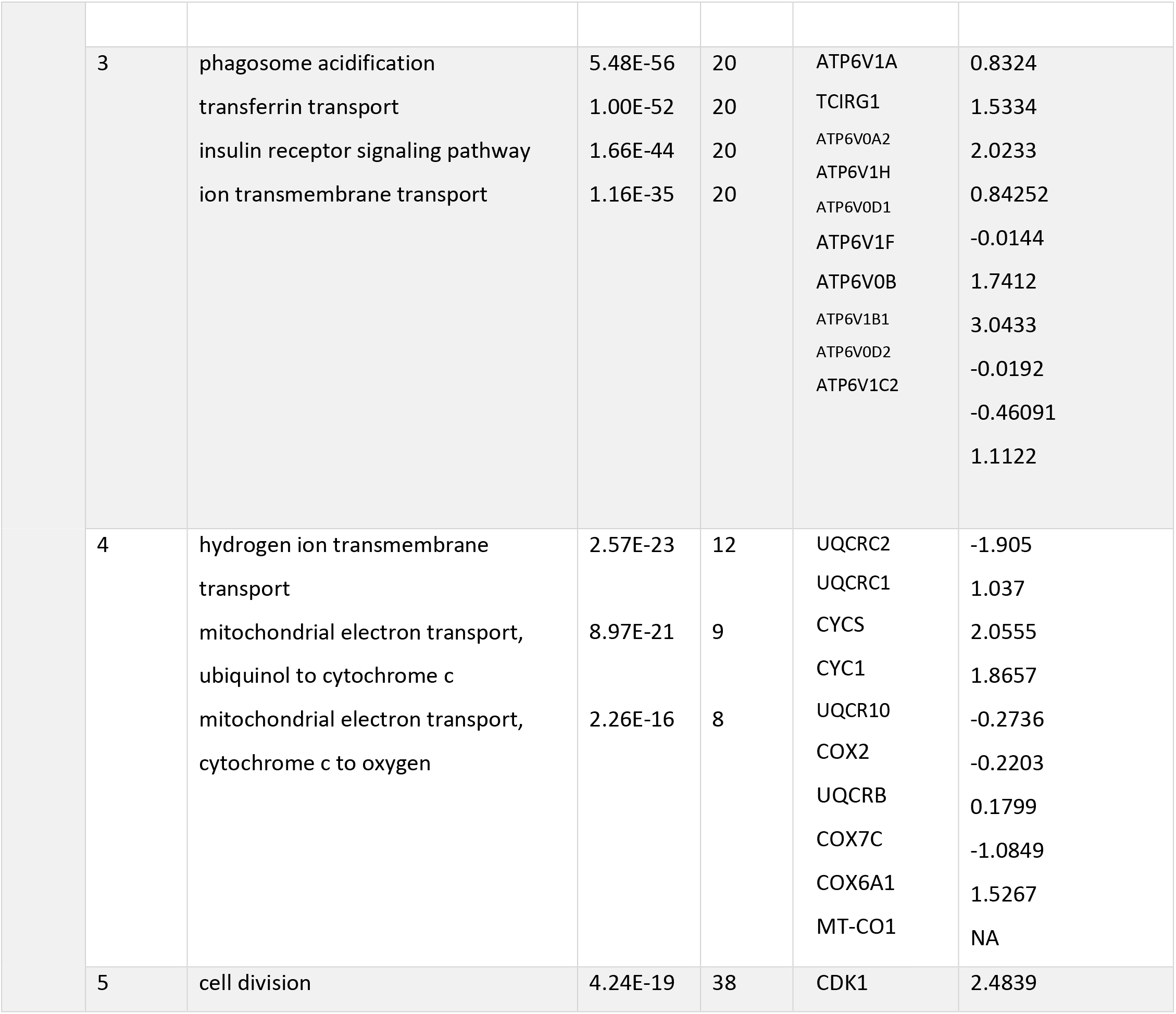
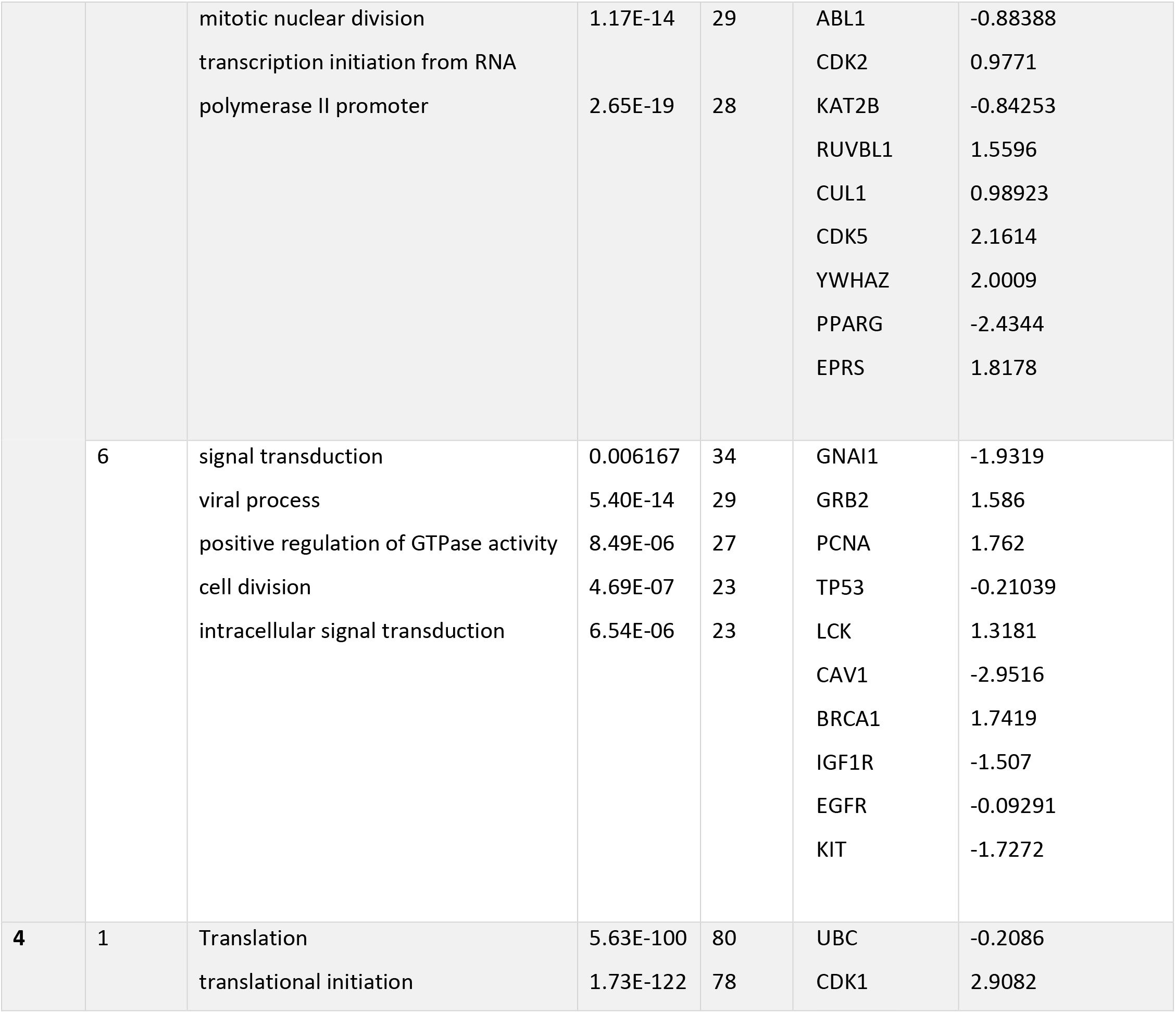
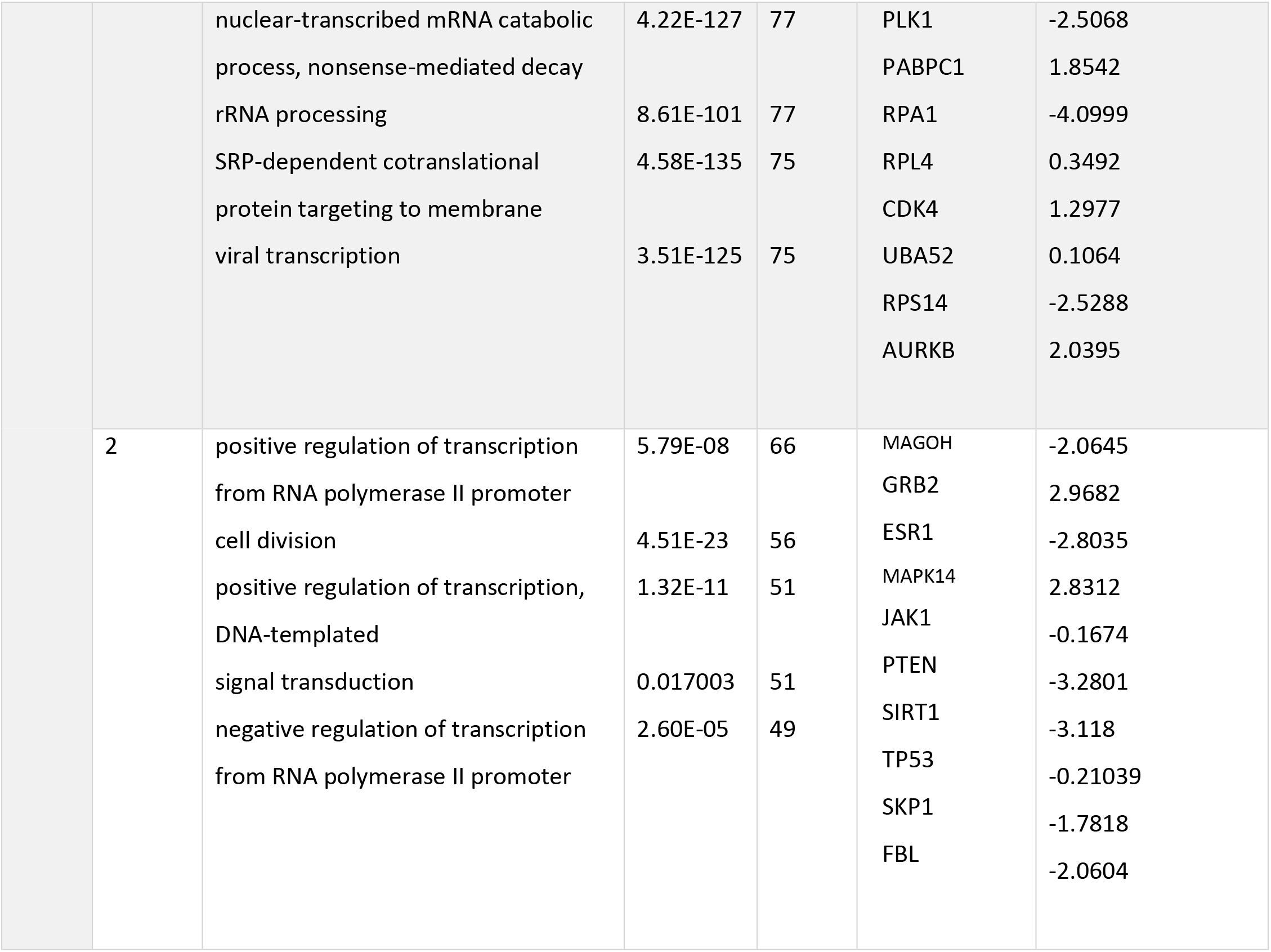
Specific Gene Ontology (GO Term), and biological process (BP) terms for each stage of breast cancer (NA: these genes are in the first order of network and are out of defined cut off). Count: number of DEGs involved in each biological process. P value: calculated by Bonferroni method.

### Prediction of the regulatory networks upstream to the DE genes recognized in the meta-analysis

To understand the upstream regulatory protein-protein interaction network of the DE genes described in the meta-analysis, a bioinformatics tool (Expression2Kinase) was used to determine: (i) the transcription factors (TFs) anticipated likely to drive the defined expression pattern; and (ii) the kinases most probably involved in the development and activation of the described regulatory complexes. Table 5 shows the top-ten predicted TFs sorted by p-value, as well as predicted kinases that could regulate the creation of these TF complexes.

**Table 5:**
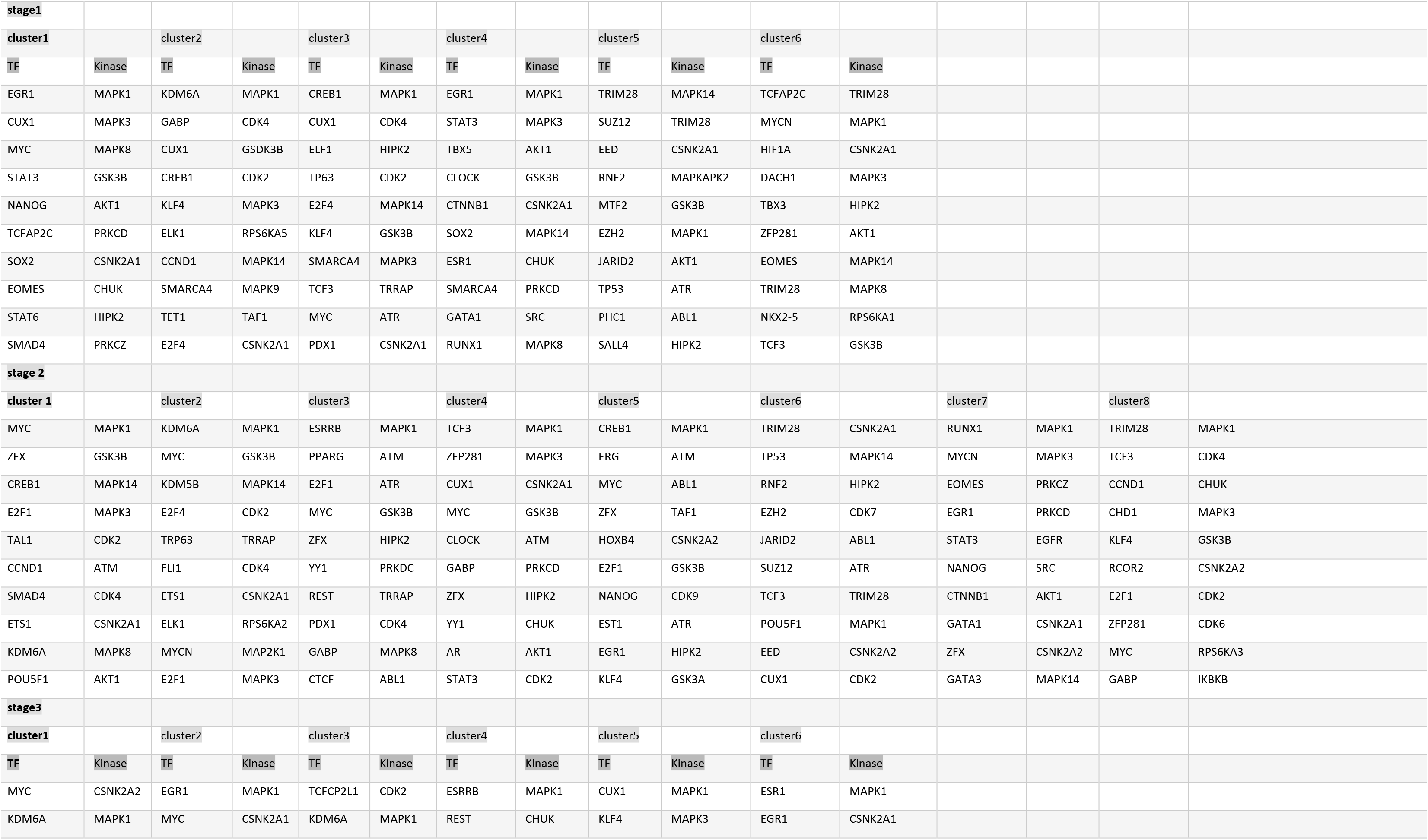
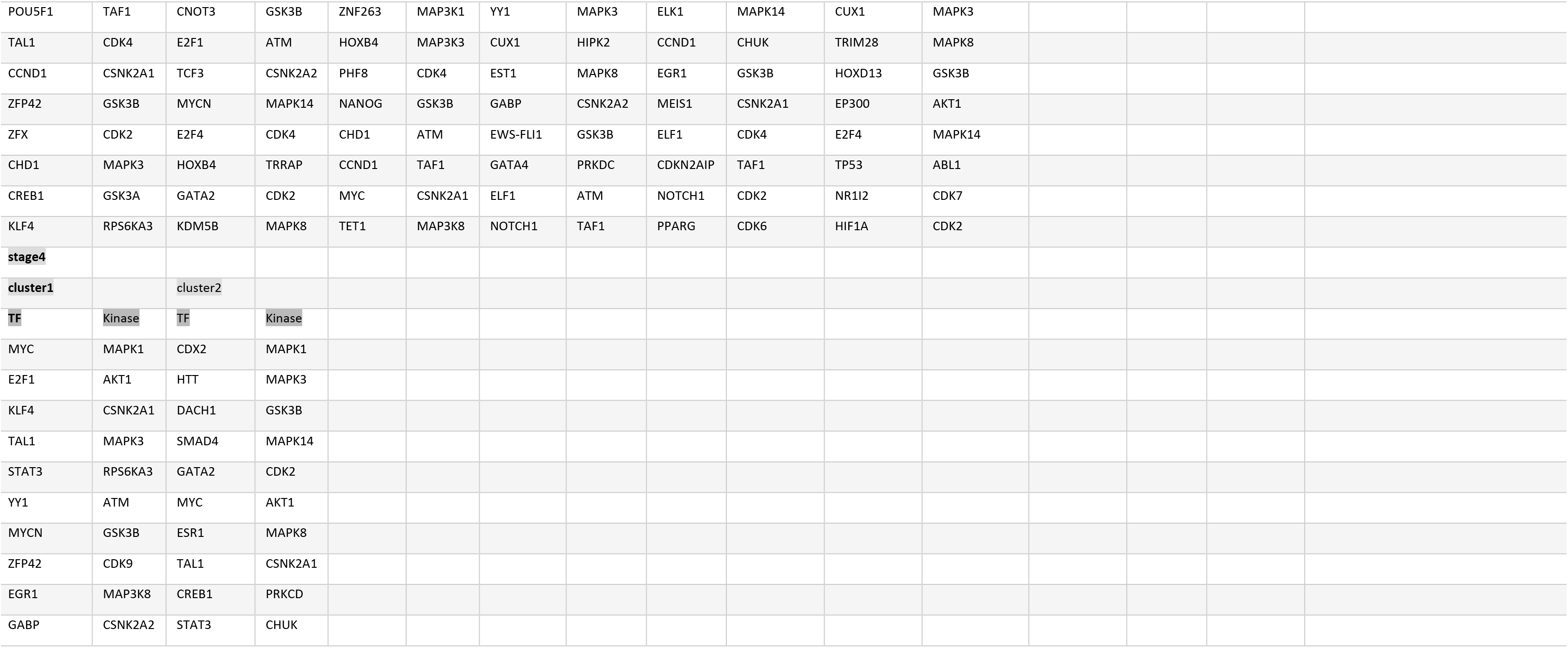
The up-stream regulatory network including transcription factors and kinases obtained from “http://www.maayanlab.net/X2K” `for four stages of breast cancer`.

## Discussion

Genome-wide screening of transcriptional alterations among normal and cancer metastases gives insight into the progression and metastasis of the molecular base of breast cancer (BC)[32]. To recognize transcriptional changes and differentially expressed genes (DEGs) in BC metastatic progression and to evaluate the prognostic function of these DEGs in clinical outcome, we assessed gene expression profiling in normal and cancer tissues using microarray technique.

The analyses of microarray data showed that many different transcripts become activated during various stages of cancer, from that, the known DEGs in this study benefit molecular detection of cancer stages.

Since, one of the main idea of this study was finding valuable biomarkers for early diagnosis and treatment, we have categorized top ten the most down and up regulated genes in each stage. In comparison with other studies, they confirmed our results about DEGs. In one study, it had reported that remarkable down-regulation of *ADAMTS5* in old patients for the first time in breast cancer[33], as well as down-regulation of this gene was observed in gastric cancer [34], but up-regulation of *ADAMTS5* was reported in colorectal[35], and lung cancer[36]. Continue on, *SCARA5* is down-regulated in breast [37, 38], and colorectal cancer[39]. In one study, *SCARA5* was reported as an immune-related molecule and could be involved in immunotherapy efforts. In fact, *SCARA5* was down-regulated due to its promoter methylation in triple negative breast cancer(TNBC) patients[38]. While, *SCARA5* overexpression significantly suppresses the proliferation of glioma cells[40]. In addition, *IGSF10* and *C2ORF40* were down-regulated [41–43]. Also, *COL10A1*, *COL11A1*, and *UHRF1* were up-regulated in breast cancer [44–46]. It is interesting that *COL11A1* was up-regulated in both stage of two and four. Actually, overexpression of miR-139-5p prevents the spread and promotes apoptosis of breast cancer cells by inhibiting *COL11A1* expression[47]. Therefore, considering its significant function and recurrence in stage two and stage four, it is useful to detect and prevent the progression of cancer in stage two. Among the top ten DEGs at all stages confirmed in this survey, *GDF10* (down-regulation) and *UBE2T* (up-regulation) were expressed only in stage two and are, in fact, stage-specific. This could be interesting for early diagnosis and treatment after further investigations using laboratory experiments. Though there are several studies that have listed these two genes in breast cancer, none of which have identified the particular stage of them. *GDF10*, a member of the TGF-β superfamily, has been reduced in tumors samples. Further study of *GDF10* expression in a wider range of clinical TNBC samples using qPCR confirmed its downregulation and association with disease severity parameters[48]. Ubiquitin-conjugating enzyme E2T (*UBE2T*) is known as up-regulated in basal-like breast tumors and amplified in breast cancer[49]. Moreover, *UBE2T* has been linked with cancer development and poor outcome of many solid tumors, including prostate or gastric cancer[50, 51]. *UBE2T* has been identified has a crucial role in the control of BRCA1 in breast cancer[52].

As an important part of this research, we aimed to detect pivotal protein hubs in PPI network with the highest betweenness index in order to gain a better insight into molecular events alongside biological pathways that play a key role in breast cancer. Furthermore, we have introduced a subset of top-ten proteins in each stage and cluster as mentioned in table 4. Among these proteins, most of them have examined in other research studies in breast cancer with down or up regulation status that showing concordance with the present study. But, we have recognized that *PLK1s1* protein in stage one was not examined in any other research studies in breast cancer. In spite of the fact that this gene is one of the top ten hub proteins with down-regulation in stage one and cluster two, it was not included in DEGs. This can be inferred that one protein can play a vital role in PPI network apart from it’s up or down regulation status. *PLK1s1* is reported as a one of the most promising genes associated with Attention-deficit hyperactivity disorder (ADHD)[53]. Another study revealed that *PLK1S1* promoted (Renal cell carcinoma) RCC tumorigenesis and enhanced sorafenib resistance of RCC through the miR‑653/CXCR5 pathway[54].

According to comprehensive analysis through preserved data in the string database, we constructed the interaction network in Networkanalyst webserver between determined DEGs and first-order neighboring nodes for each stage to have broader vision over altered cellular events, and consequently ascertain key targets, better concept of treatment in association with each stage of cancer development.

The main network was split into smaller topological segments for more accurate view and further complimentary assessment on whole created networks and each subnetwork. Each subnetwork is composed of various functional parts which depicted the perfect overview of molecular changes, occurred in breast cancer [55](Table 4) .The molecular changes mostly include an increase in genes expression at the translational level, enhanced regulatory processed of cell division and segregation of daughter chromatids. Wnt signaling is one of the most important pathways in cancer development, which trigger breast tissue remodeling during gestation and milking periods [56, 57]. This pathway shows more activation at stage one, but there has been no report of the presence of Wnt signaling related protein in tumor cells [58]. The results of this study along with reported expression change for some of the mainly involved proteins in this pathway such as WNT10A and WNT6 show up-regulation [59–62], while WNT11 displays down-regulation which is at odds with Dwyer and colleagues’ studies [63].

At stage two, on one hand, due to an increase in cell energy demand, the aerobic mechanism of energy creation like mitochondrial function and electron transfer chain became activated. On the other hand, anaerobic respiratory pathways would start, in other words, catabolic process related to anaphase inducer complex, proteasome related, and protein destruction pathway were activated through ubiquitin ligase activity, leading mitotic cell cycle, particularly in tumor condition. Studies of Hugute, Vouyovitch, and their colleagues reported the increased presence of other proteins from WNT signaling such as WNT4 at this stage, but in our survey this protein show down-regulation [64, 65]. We noticed that expression of WNT7B show increased that could be the outcome of altered cellular requirements. In another study, Hugute et al undeniably revealed that WNT2, WNT4, WNT7B signaling proteins play key roles in abnormal cell division in breast cancer. But WNT10A protein shows a decrease in its expression at the second stage, compared to the first one,which could indicate that cells are moving towards a lower level of differentiation and evolution [66, 67]. WNT11 protein shows down-regulation at stages one and two (KEGG=hasa04310), could provide an opportunity for cell detachment from its niche. This protein triggers a focal connection through Planar Cell Polarity (PCP) pathway. In this direction, it has been reported that positive regulatory pathways of small GTPase also become activated. The various study showed that small GTPase affects various cancer occurrence [68–71]. As inhibition of these pathways would lead to obstruction of cancer development towards upper stages and prevent metastasis, thus more investigations have been made to detect the key proteins such as MAGOHB. At the 2nd stage of breast cancer, the expression of MAGOHB is increased, as visualized in cluster two of the constructed network. Analyses showed that removal of this protein and its paralogue would reduce cell survival [72]. This protein, however, is associated with increased expression in the three early stages, which leads to increased survival of tumor cells and can, therefore, be identified as one of the potential therapeutic targets in breast cancer. To sum up, stage two refers to the changes that make the cell preparation for a series of fundamental changes.

In addition to translational alterations, transcription, replication, increased mitochondrial and electron transfer chain pathways to generate more energy, modifications such as acidification of phagosomes as phagocyte degradation sites and even alterations of one of the cell surface receptors have been well studied in breast cancer cell lines. Moreover, it has been clearly shown that increased activity of this pathway is directly correlated with the metastatic levels of breast cancer cells [73–76].

In the study of Shen et al, increased expression of the ferritin transmission system suggesting a different degree of cell need, since at this point the cells require large amounts of energy[77]. On the other hand, due to changes in the expression of its receptors, the cytoskeleton lead to contact-inhibition that will be less closely related to the environment and ready to be separated from tumor and basal tissue. Moreover, in step 4, it migrates to other tissues that are more closely related to molecular markers. WNT9a is a separate family member of WNT, and research indicates that this protein could have a beneficial impact on cell proliferation and differentiation via IHH gene expression. In addition, as clearly stated in the Saptter el al report, IHH protein is one of the key factors involved in skeletonogenesis, which could lead mainly metastasis of the breast tissue [77]. On the other hand, Irshad ali et al confirmed that a twenty three-times increase in WNT9a protein expression followed by a decrease in colorectal cell proliferation, and their analysis showed that this protein had a two-times increase in expression compared to control (based on log2 unit). [78]

All aforementioned suggested that some kind of changes occurred in breast cells, which eventually end up with tumor condition and metastasis. But none of the initial alterations at the commence of cancer development are definite indicators of cancer occurrence, in fact, each change is originated from and closely associated with its upstream one(s) in related pathways. Thus, we investigated the upper level of differential expressed genes (DEGs) at regulatory networks in which transcription factors and protein kinases are functioning. (Table5).

Many of these TFs and kinases are involved in different clusters at the first stage and some are specific to each cluster. Investigations showed that these proteins or factors are highly effective concerning cancer occurrence [79]. Studies on these proteins revealed their profound effect on breast cancer emergence. For example, the presence of EGFR1 protein in cluster 1 and 4 showed down-regulation in breast cancer and consequently its tumor inhibitory effect displayed reduction. Moreover, it is clearly shown that increases in the expression of this protein trigger inhibition of cancer cell proliferation [80], while, the opposite results were reported in prostate cancer [81]. And this protein increases the drug resistance through an increase in MDR1 expression in breast cancer cells [82], resulting prevention of drug effects on these cells. Also, studies on upstream regulatory parts of EGR1depicted that this region is composed of several sections which could play a pivotal role in the present level of this protein inside cells [83–85]. Further investigations are required to elaborate upon more on the alteration of the promoter region of this gene in breast cancer.

Furthermore, the transcription factor CUX1 which is the key protein of three initial clusters has various functions in association with the sequence of effective genes promoters, which would account for an investigation into the promoter region of those genes that are regulated by this protein [86]. Sansegeret et al researches clearly showed that increase in time length and level of CUX1 presence triggers chromosomal instability in breast cancer cells [87]. In agreement with their study, we noticed the enhanced expression of CUX1 gene, which could be considered as one of the factors, leading cell towards tumor status. Evaluation of this protein function, using COSMIC database (https://cancer.sanger.ac.uk/cosmic/census-page/) illustrates that this protein plays a role on various pathways or biological functions, triggering tumor statuses such as apoptosis, immune system avoidance, vascularization, invasion, metastasis, and developmental processes. Also, SOX2 is another important transcription factor (TF) whose oncogenic role on cancer occurrence has been perfectly studied [88]. However, the expression level of this gene in our analyses was lower than the appointed threshold log<2 (Table5). These proteins are specific for each stage and are valuable for independent study.

It has been known that MAPK1 is one of the key kinases in various cancers development that was identified in this study. Similarly, recent studies show the effect of MAPK1 protein on miR-585 expression concerning gastric cancer occurrence [89]. In this analysis, the expression of this gene was displayed less than appointed threshold and consequently, MAPK1 was not listed among DEGs. Another active protein kinase in tumor condition is MAPK3 and according to protein atlas database (https://www.proteinatlas.org/), the level of this protein presence is more than the expression of its corresponding mRNA, this could point out the longer half-life of transcription in most of the soft tissues including adipose tissues of the breast. Alteration in mRNA half-life in these tissues may be one of the reasons that would increase the chance of cell proliferation and tumor emergence [90, 91].

The already discussed issues imply the importance of the regulatory up-stream network of the identified DEGs in this study. Further investigation into the studied protein kinases and transcription factors, could lead to the detection of protein kinases in all stages of breast cancer development, including AKT1. Anvar et al showed the interaction between AKT1 and other proteins, involved in gastric cancer cells. This cross-talk may has been identified in breast cancer, as well [92]. One interesting point concerning stage one is that the significant increase in transcription factors (TFs) relative to cluster five is unique and non-repetitive compared to other TFs active at this stage. Thus, probably, at least two different pathways are functional in stage one of breast cancer. However, it is not clear whether this network of transcription factors is independent of other transcription factors. From that, we constructed the interaction network between TFs of cluster five and other TFs, obtained in GeneMANIA[93]. The outcome shows that TRIM28 has the most betweenness and closeness indices (P-value<0.05 and logarithmic Fold Change (logFC=1.78). Li et al studies showed that TRIM28 protein in association with EZH2 (histone methyl transferases) is expected to not only co-regulate the inhibitory effect on genes expression but also activate transcription, resulting from cancer development [94]. In addition, the research studies of Czerviska and colleagues has unequivocally shown that an increase in TRIM28 protein causes an increase in the invasion and development of tumor stem cells in breast cancer. [95]. Oleksiewicz et al researches have shown that TRIM28 protein along with KRAB-ZNFs pathway could trigger an alteration in DNA methylation pattern [96]. Therefore, perhaps one of the pathways resulting in breast cancer development could start with changes in the methylation pattern of genes or alteration in the expression of genes, involved in the modification of DNA methylation pattern. TRIM28 can be one of the key targets for treatment through methylation, whereas the expression modification in most of these genes are lower than the appointed threshold level.

Based on the above, it can be concluded that breast cancer is likely to have a modification in the methylation pattern of DNA in the regulatory regions of some genes, such as TRIM28, which leads to other expression changes through the interaction between TRIM28 affected proteins.

The investigation into an involved regulatory phase in extracted clusters of constructed networks has revealed that each stage of cancer development benefits from its specific TF and kinases, which would be recognized as biomarkers as well as potential diagnosis targets for cell proliferation termination in breast cancer. Some of these key proteins are STAT6 ،GSDK3B ،RPS6KA5 ،MAPK9 ،TP63 ، TBX5 ،MTF2 ،PHC1 ،SALL4 ،MAPKAPK2 ،TBX3 and NKX2-5, RPS6KA1 (stage1), RCOR2 ،EGFR ،GATA3 ،ERG ،AR ،CTCF ، MAPK2K1 ،RPS6KA2 ،FLI1, TRP63 (stage2), CNOT3 ،TCFCP2L1 ،ZNF263 ،PHF8 ،MAP3K1 ،MAP3K3 ،EWS-FLI1 ،GATA4 ، MEIS1 ،CDKN2AIP, HOXD13 ،EP300, NR1I2 (stage3), and CDX2 و HTT (stage4). In other words, these proteins obtained from X2K analysis and each of which can be study separately for the future.

In conclusion, the suggested keynote of breast cancer studies is that every gene expressed is not necessarily translated, and some advanced and meticulous mechanism has a role during the whole process. Therefore, we recommend studying gene expression and protein presence simultaneously [97].

